# Circulating biomarkers in serum from aortic valve stenosis patients predict sex-specific drug responses in valve myofibroblasts

**DOI:** 10.64898/2026.01.16.700010

**Authors:** Brandon J. Vogt, Megan Chavez, Nicole E. Félix Vélez, Rayyan M. Gorashi, Ryan R. Reeves, Brian A. Aguado

## Abstract

Aortic valve stenosis (AVS) is a prevalent, sexually dimorphic cardiovascular disease characterized by fibro-calcification of the aortic valve leaflet. Sex differences in AVS arise in part from sexually dimorphic serum composition that differentially regulate valvular interstitial cell (VIC) myofibroblast activation. However, how individual serum factors contribute to sex-specific drug responses targeting VIC myofibroblast activation remains unknown. Here, we integrate serum proteomic profiling with *in vitro* drug screening using hydrogel biomaterials to identify sex-specific regulators of antifibrotic drug efficacy. We found that Insulin-like Growth Factor Binding Protein 2 (IGFBP2) serum levels are associated with resistance to the antifibrotic drug Evogliptin only in female VICs cultured with female AVS serum. This mechanism is driven by IGFBP2-mediated activation of Rho/ROCK and focal adhesion kinase signaling pathways that counteract Evogliptin treatment. Our findings reveal a sex-specific, serum-mediated mechanism of Evogliptin resistance and highlight IGFBP2 as a candidate biomarker for stratifying female AVS patients for Evogliptin treatment. More broadly, these findings underscore the importance of incorporating sex-stratified biomarker analyses into AVS therapeutic development to improve patient-specific treatment recommendations.

## Introduction

Aortic valve stenosis (AVS) is characterized by extensive fibrosis and calcification of the aortic valve leaflet that impedes cardiovascular function.^1,2^ As AVS develops, patients experience impaired hemodynamics, left ventricular pressure overload, and eventual heart failure.^3^ AVS risk increases dramatically with age and is highly prevalent in elderly populations with approximately 1 in 8 adults over the age of 75 affected.^4^ Moreover, population aging is expected to drive increases in both AVS incidence and treatment costs, with an estimated 19 million individuals expected to be affected by 2036 and costs projected to reach $19 billion by 2034.^5–7^ Currently, AVS is treated through interventional procedures such as surgical or transcatheter aortic valve replacements which implant a new valve to restore normal hemodynamic function in the heart and prolong patient survival.^8,9^ However, these interventions pose their own procedural risks and only serve as temporary solutions as patients can experience failure of the newly implanted valve.^10,11^

To address these limitations, researchers and clinicians have long pursued a variety of pharmacologic strategies to slow or halt AVS progression, resulting in multiple clinical trials.^12–16^ Statins are the most extensively investigated drugs for AVS, owing to their proven efficacy in atherosclerosis, which involves similar pathogenic processes such as lipid accumulation and oxidation.^17,18^ However, large-scale trials of statins and other lipid-lowering drugs have yielded little to no success in halting AVS progression.^14–16^ Other drug classes, including anti-inflammatory drugs and bisphosphonates have also been explored with similar disappointing outcomes.^12,13^ More recently, novel inhibitors such as Evogliptin have shown early promise in suppressing valvular calcification.^19–21^ However, despite these efforts, current therapeutic options remain largely limited to symptom management, including antihypertensive and renin-angiotensin-aldosterone system-targeting drugs to reduce cardiac load.^22^ Given there are still no approved drugs to reverse AVS progression, there is an active need for patient-specific treatment strategies.

AVS is a heterogeneous disease with significant patient-to-patient variability and a wide range of disease phenotypes, likely necessitating a precision medicine approach for identifying effective drug treatments.^23–25^ Specifically, AVS progression is sex-dependent, with male AVS patients experiencing elevated calcification of the aortic valve relative to female AVS patients who instead show increased fibrosis at the early stages of disease.^26–28^ Observed sex differences in AVS progression extend to clinical treatment responses such as increased mortality following surgical valve replacement in female patients and increased mortality following transcatheter valve replacement in male patients.^29–31^ Sex-based disparities are further compounded by the fact that female AVS patients are more often underdiagnosed and are referred for intervention at later disease stages than men.^32–35^ As a result, researchers and clinicians have developed sex-specific risk profiles for the recommendation of valve replacement procedures.^36,37^

Interestingly, sex differences in AVS have also been linked to the biological sex of aortic valve cells. Male and female valvular interstitial cells (VICs), the resident fibroblast of the aortic valve, show divergent phenotypes in response to mechanical and biochemical cues.^38,39^ These sex differences are best captured on hydrogel biomaterials used as cell culture platforms that recapitulate the stiffness of aortic valve tissue.^40–42^ Specifically, female VICs with XX chromosomes cultured on hydrogels show elevated myofibroblast activation, a pro-fibrotic phenotype associated with extracellular matrix turnover and collagen deposition.^38,43^ In contrast, male VICs with XY chromosomes cultured on hydrogels are more prone to differentiating into an osteoblast-like phenotype that promotes calcification of the aortic valve.^44,45^ Cellular scale sex differences also regulate drug responses, with recent findings showing that sex chromosome linked genes contribute to sex-specific drug efficacy in VICs.^38^ These differences have been leveraged to develop sex-specific predictive models of VIC drug responses and identify sex-biased drug treatment parameters to improve drug responses in both male and female VICs.^46^ Collectively, these findings suggest that sexually dimorphic VIC phenotypes play a central role in AVS progression and may contribute to variations in clinical treatment outcomes.

Inflammatory factors present in AVS serum have also been shown to promote pathological VIC phenotypes.^41,47^ AVS serum contains a myriad of cytokines, growth factors, and extracellular matrix components that can individually and collectively interact to alter VIC function.^48,49^ Notably, following transcatheter aortic valve replacement, the serum proteome undergoes significant changes that are broadly associated with maintaining VIC quiescence.^50^ Interestingly, serum samples from male and female patients show enrichment of distinct clusters associated with AVS progression such as increased actin binding in female serum and increased cholesterol metabolism in male serum.^51^ Moreover, patient-specific serum proteomic profiles have been used to accurately predict VIC drug responses.^52^ These findings suggest that the serum proteome contributes to patient heterogeneity, individualized drug responses, and sex-specific mechanisms of AVS development. However, the mechanisms by which individual serum factors influence VIC drug responses in a patient-and sex-specific manner remain unknown.

In this study, we utilized hydrogel biomaterials and serum proteomic analyses to identify predictive links between serum factor abundance and myofibroblast responses to antifibrotic drugs *in vitro*. We hypothesized that individual serum biomarkers would be predictive of drug responses in VICs and that male and female serum contain distinct biomarkers that contribute to patient-specific drug sensitivity. To test these hypotheses, we cultured male and female VICs with sex-matched AVS patient serum and treated them with one of three antifibrotic drugs that are in ongoing clinical trials for treating AVS: Ataciguat (NCT07001800), Evogliptin (NCT05143177), or Colchicine (NCT05162742).^19–21^ By correlating *in vitro* drug responses with serum protein abundance, we identified a female-specific association between elevated insulin-like growth factor binding protein 2 (IGFBP2) levels and reduced efficacy of the dipeptidyl peptidase 4 inhibitor, Evogliptin. Through RNA sequencing, we found that female serum IGFBP2 levels regulate Evogliptin sensitivity in VICs through Rho/ROCK and focal adhesion kinase (FAK) signaling pathways. These findings were confirmed through neutralizing antibody experiments to block the effects of IGFBP2 and inhibiting Rho/ROCK and FAK signaling to restore Evogliptin efficacy in female VICs cultured with female serum. Together, these findings highlight IGFBP2 as a key mediator of Evogliptin resistance in female VICs and a potential biomarker for patient stratification in ongoing clinical trials.

## Results

### Proteomic profiling reveals distinct serum signatures in AVS patients

We first sought to identify proteins that are upregulated in AVS patient serum relative to healthy controls. Based on echocardiographic and patient data, the male AVS cohort was younger and had a 1.5-fold larger aortic valve area relative to the female AVS cohort (**Table S1**). All serum samples were analyzed using the SomaLogic DNA aptamer–based proteomic platform^53,54^, which quantified the relative abundance of a pre-selected panel of 1,512 proteins with known associations with inflammation or cardiovascular disease. To increase the size of our healthy control group, we supplemented our cohort with 38 age-matched individuals from the SomaLogic serum proteome database (**Figure 1A**). Unsupervised hierarchical clustering revealed that AVS and healthy serum samples largely cluster together, suggesting significant proteomic changes occur during AVS progression (**Figure 1B**). Principal component analysis confirmed that healthy serum samples cluster separately from AVS serum and that AVS serum exhibits markedly greater heterogeneity (**Figure 1C**).

**Figure 1.**
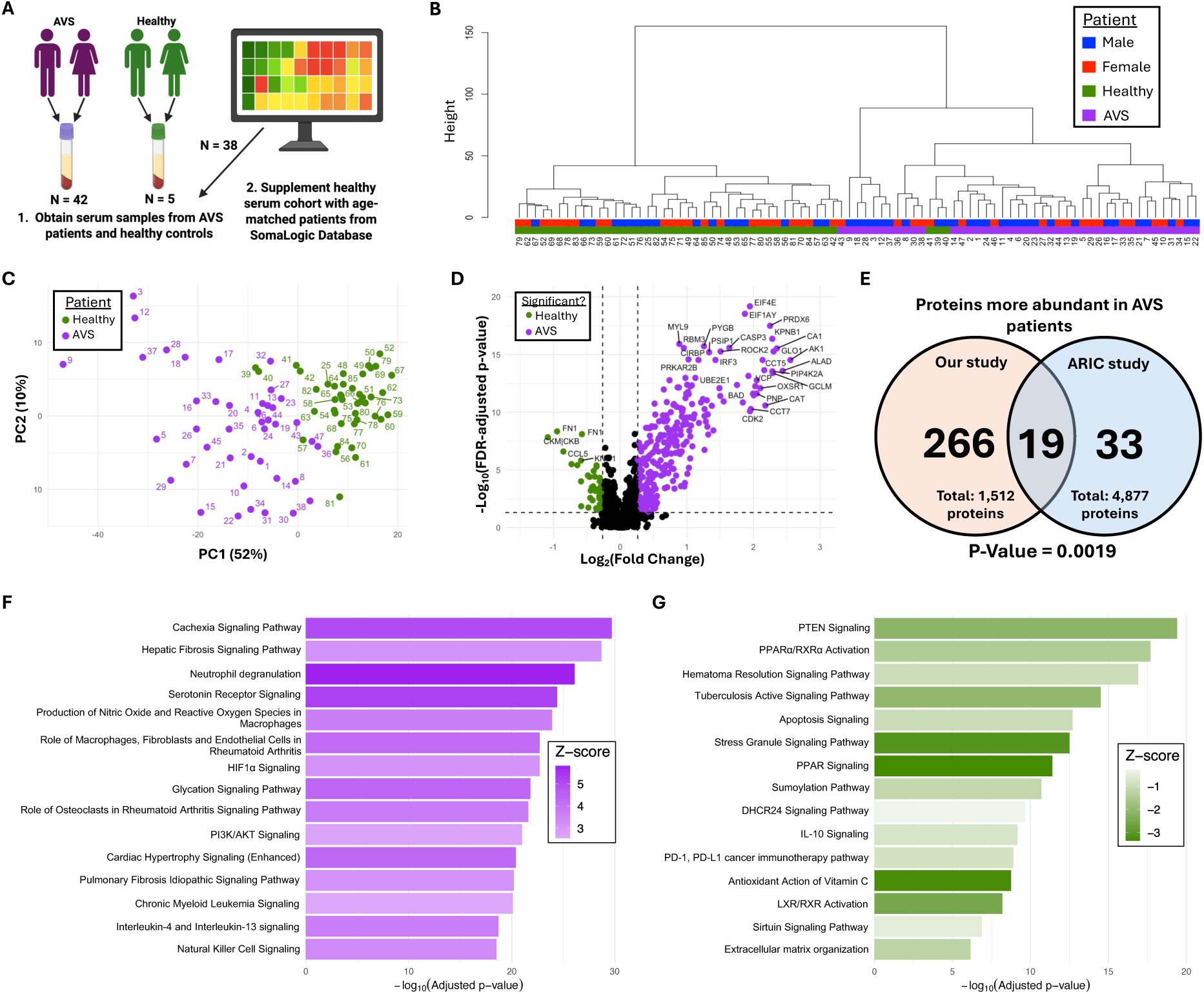
Proteomic analysis of AVS patient serum. A, Schematic overview of the approach to obtain and supplement serum samples for proteomic analysis. **B,** Hierarchical clustering dendrogram linking all serum samples based on proteomic data (N=43 healthy, N=42 AVS). **C,** Principal component analysis of the serum proteome from healthy and AVS patients (N=43 healthy, N=42 AVS). **D,** Volcano plot of 1,512 protein abundances in healthy serum samples (green) or AVS serum samples (purple) (N=43 healthy, N=42 AVS). Statistical significance was defined as false-discovery rate (FDR) adjusted P-value < 0.05 (unpaired t-test with Welch’s correction) and fold change greater than 1.2. **E,** Venn diagram comparing serum biomarkers identified in this study against the serum biomarkers identified in the Atherosclerosis Risk In Communities study.^55^ Statistical significance was determined by Fisher’s exact test. **F** through **G,** Ingenuity Pathway Analysis showing the top pathways associated with proteins that are more abundant in **(F)** AVS serum samples and **(G)** healthy serum samples.

We next performed a differential protein analysis and identified 41 proteins more abundant in healthy serum and 285 proteins more abundant in AVS serum (**Figure 1D**). A similar comparison between male and female AVS serum revealed minimal baseline sex differences, with only five proteins more abundant in female AVS serum, and none more abundant in male AVS serum (**Figure S1**). To further narrow down candidate AVS biomarkers, we compared our identified proteins with AVS biomarkers identified from the Atherosclerosis Risk In Communities study that tracked over 11,000 patients over three decades.^55^ We found a significant overlap between the two data sets, corroborating our findings and narrowing down our biomarker pool to 19 robustly validated serum proteins (**Figure 1E**). Using all 285 serum factors more abundant in AVS patient serum, we used Ingenuity Pathway Analysis (IPA) to identify pathways altered during AVS progression. The top enriched pathways were linked to fibrotic and cardiovascular disease processes, inflammatory signaling, and altered immune cell function (**Figure 1F**). Complementary analysis using the Kyoto Encyclopedia of Genes and Genomes (KEGG) database revealed similar results, with the top pathways linked to inflammation and cardiovascular disease (**Figure S2**). We also used IPA on proteins more abundant in healthy serum and observed enrichment of anti-inflammatory and metabolic pathways (**Figure 1G**). However, KEGG analysis did not identify any significantly enriched pathways, likely due to the limited number of proteins upregulated in the healthy cohort.

### Clinically relevant doses of Ataciguat, Colchicine, and Evogliptin reduce myofibroblast activation in male and female VICs

Next, we optimized doses of three antifibrotic drugs currently in clinical trials for treating AVS (Ataciguat, Colchicine, and Evogliptin).^19–21^ Male or female VICs were treated with fetal bovine serum (FBS) and either 5%, 10%, or 50% of each drug’s maximum serum concentration (Cmax) (**Table S2**). After two days of treatment, VICs were immunostained for alpha smooth muscle actin (αSMA) and cleaved caspase-3 to assess myofibroblast activation and cell viability (**Figure 2A**). All experiments were performed using our previously optimized poly(ethylene glycol) hydrogel formulation with an elastic modulus of ∼51 kPa (**Figure 2B**).^56^ This platform mirrors the stiffness of diseased aortic valve tissue and provides a physiologically relevant cell culture environment known to influence sex-specific myofibroblast activation and drug sensitivity.^46^

**Figure 2.**
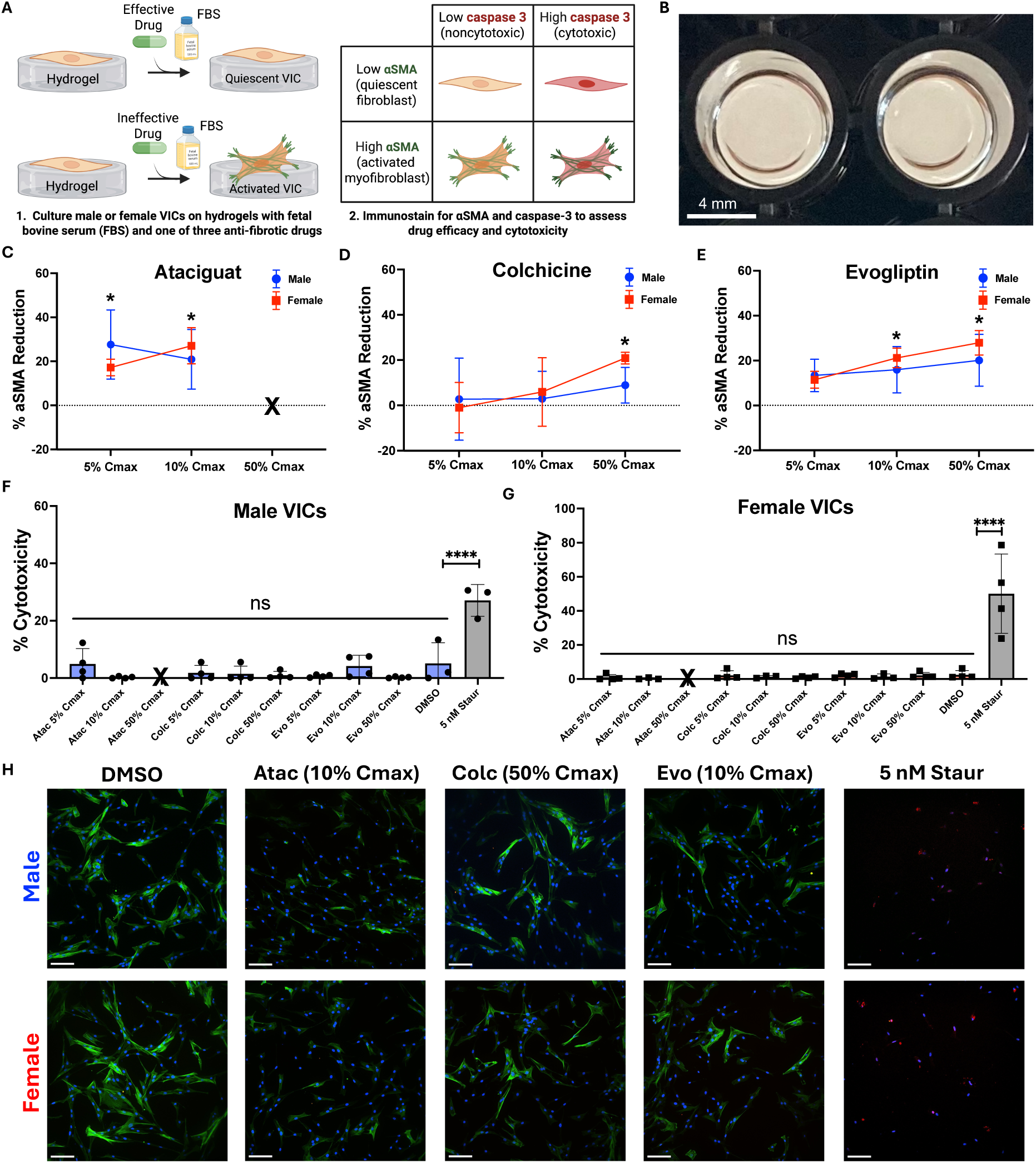
Optimizing clinically relevant doses of antifibrotic drugs. A, Schematic overview of the protocol to optimize antifibrotic drug dosing using hydrogel biomaterials. **B,** Representative image of formed hydrogels ready for cell seeding. **C** through **E,** Percent αSMA reduction in male and female VICs cultured on hydrogels with increasing Cmax concentrations of **(C)** Ataciguat, **(D)** Colchicine, or **(E)** Evogliptin (N=3-4 gels). X indicates complete cell death at that concentration. Statistical significance is indicated as * = P < 0.05 (one sample t and Wilcoxon test against a theoretical mean of zero). **F** through **G,** Percent cytotoxicity in **(F)** male and **(G)** female VICs in response to increasing Cmax concentrations of Ataciguat, Colchicine, or Evogliptin (N=3-4 gels). Statistical significance is indicated as **** = P < 0.0001 (one-way ANOVA with Tukey posttests). **H,** Representative immunofluorescent images of male and female VICs cultured on hydrogels with different experimental conditions (green: αSMA, blue: nuclei, red: cleaved caspase-3). Scale bar = 100 μm. Abbreviations: Atac = Ataciguat, Colc = Colchicine, Evo = Evogliptin, Staur = Staurosporine.

Using these hydrogels, we found that Ataciguat had a maximal efficacy at a 10% Cmax dose, as indicated by the greatest reduction in αSMA gradient mean intensity relative to vehicle control (**Figure 2C**) Ataciguat at a 50% Cmax dose was fully cytotoxic and not quantified. Colchicine and Evogliptin were similarly effective in male and female VICs at 50% and 10% Cmax doses, respectively (**Figure 2D-E**). No sex differences in drug efficacy were observed across any dosing parameters tested when using fetal bovine serum. Importantly, all optimized drug doses were confirmed to have minimal cytotoxicity in both male and female VICs, suggesting that optimized drug concentrations are both efficacious and well-tolerated (**Figure 2F-H**).

### Sex-specific serum biomarkers predict antifibrotic drug efficacy

We next sought to determine biomarkers in human serum that regulate antifibrotic drug efficacy *in vitro*. To test this, male or female VICs were seeded on hydrogels and treated with sex-matched serum and optimized doses of Ataciguat, Colchicine, or Evogliptin. After two days, VICs were fixed and immunostained for αSMA to quantify myofibroblast responses to each drug. For each serum sample, drug efficacy was plotted against the relative abundance of each of the 19 candidate biomarkers to generate robust, sex-specific correlations (**Figure 3A**). At baseline, female VICs cultured with female AVS serum exhibited a 1.4-fold higher rate of αSMA expression relative to male VICs cultured with male AVS serum (**Figure 3B**). Additionally, Ataciguat demonstrated a 1.6-fold greater efficacy in reducing myofibroblast activation in female VICs relative to male VICs, whereas Colchicine and Evogliptin were similarly effective in both sexes (**Figure 3C**).

**Figure 3.**
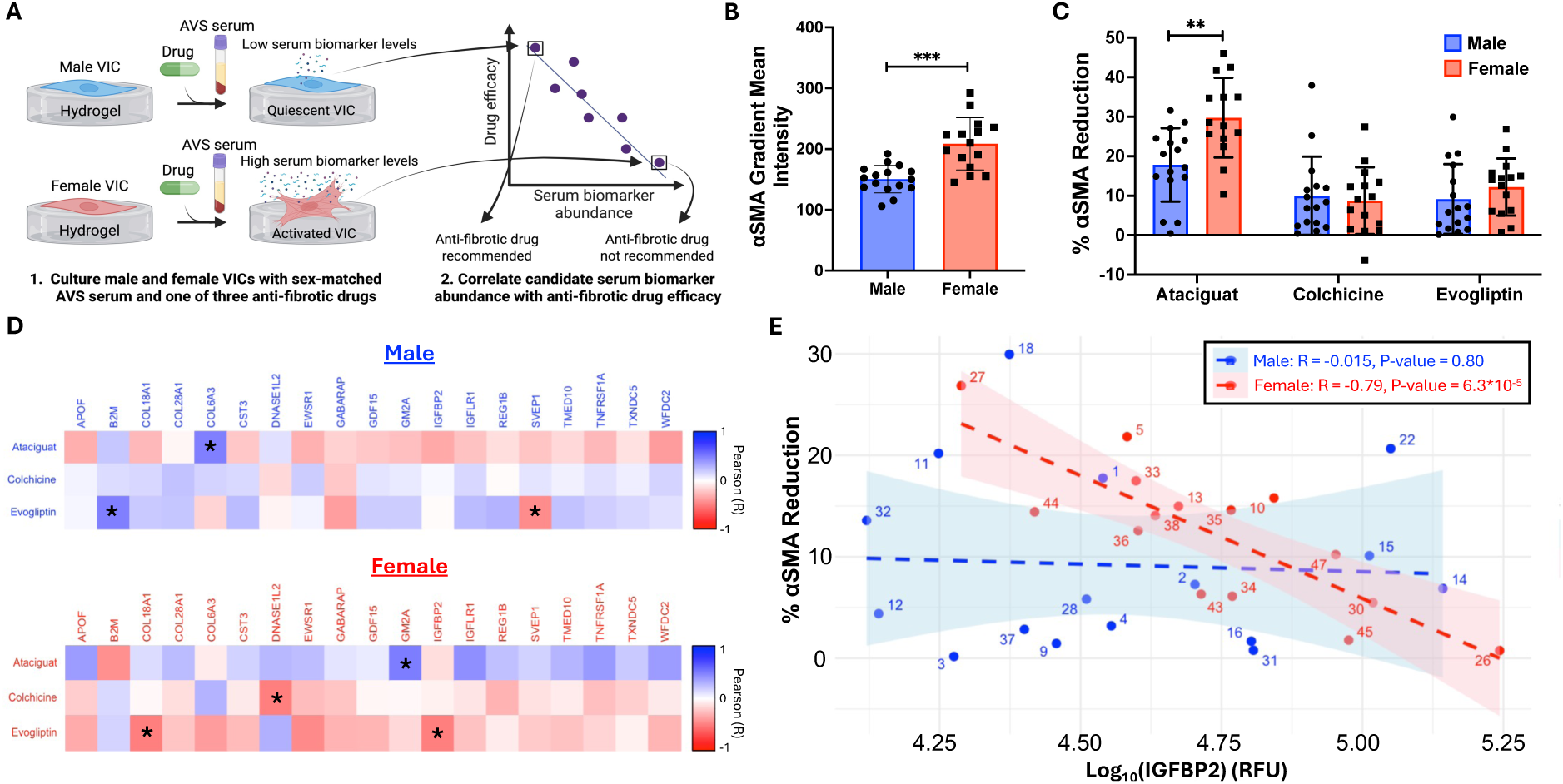
Identifying serum factors that regulate antifibrotic drug efficacy. A, Schematic overview of the approach to link candidate serum biomarkers with *in vitro* drug efficacy. **B,** αSMA gradient mean intensity for male and female VICs cultured on hydrogels with sex-matched AVS patient serum (N=16 male, N=15 female). Statistical significance is indicated as *** = P < 0.001 (unpaired two-tailed t-test with Welch’s correction). **C,** Percent αSMA reduction in male and female VICs cultured on hydrogels with optimized doses of Ataciguat, Colchicine, or Evogliptin. Statistical significance is indicated as ** = P < 0.01 (unpaired two-tailed t-test with Welch’s correction). **D,** Heatmaps showing the correlation between each candidate serum factor and antifibrotic drug efficacy for male or female VICs cultured with sex-matched AVS serum (N=16 male, N=15 female). Statistical significance is indicated as * = P < 0.05 (paired two-tailed t-test). **E,** Correlation between serum IGFBP2 abundance (relative fluorescence units (RFU)) and percent αSMA reduction in male and female VICs cultured on gels with sex-matched AVS serum and Evogliptin (N=16 male, N=15 female).

Overall correlations between drug efficacy and serum factor abundance were first assessed by pooling male and female samples (**Table S3)**. Across all samples, we identified three significant correlations between serum protein abundance and Evogliptin efficacy, whereas no significant correlations were observed with Ataciguat or Colchicine (**Figure S3**). Sex-specific analyses revealed that correlations between drug efficacy and serum biomarker abundance differed significantly between male and female samples (**Tables S4 and S5**). More specifically, female VICs cultured with female AVS serum exhibited predominantly positive correlations between serum biomarker abundance and Ataciguat efficacy, and predominantly negative correlations for Colchicine or Evogliptin. In contrast, male VICs cultured with male AVS serum displayed the opposite trend with serum factor abundance negatively correlating with Ataciguat efficacy and positively correlating with Colchicine and Evogliptin efficacy (**Figure S4**).

Next, we determined male-specific and female-specific correlations between drug efficacy and individual serum factor abundance. In male VICs cultured with male AVS serum, Collagen alpha-3(VI) chain (COL6A3) correlated positively with Ataciguat response (Pearson’s R = 0.495), and Beta-2-microglobulin (B2M) correlated positively with Evogliptin (Pearson’s R = 0.497), whereas Sushi, von Willebrand factor type A, epidermal growth factor, and pentraxin domain containing 1 (SVEP1) showed a significant negative correlation with Evogliptin (Pearson’s R =-0.444) (**Figure 3D**). In contrast, in female VICs cultured with female AVS serum, Ganglioside GM2 activator (GM2A) was positively correlated with Ataciguat efficacy (Pearson’s R = 0.510), while Deoxyribonuclease-1-like 2 (DNASE1L2) was negatively correlated with Colchicine efficacy (Pearson’s R =-0.531). Additionally, Collagen alpha-1(XVIII) chain (COL18A1) and Insulin-like growth factor binding protein 2 (IGFBP2) were negatively correlated with Evogliptin efficacy (Pearson’s R =-0.613, Pearson’s R =-0.791). Of these correlations, the strongest sex-specific association was the female-specific negative correlation between serum IGFBP2 abundance and Evogliptin efficacy (**Figure 3E**). Interestingly, male serum IGFBP2 levels did not correlate with Evogliptin response, suggesting that IGFBP2 may uniquely regulate Evogliptin sensitivity in female VICs cultured with female AVS serum.

### IGFBP2 promotes Evogliptin resistance only in female VICs cultured with female AVS serum

We next tested whether IGFBP2 can directly modulate female VIC responses to Evogliptin. First, we observed that IGFBP2 is more abundant in AVS serum relative to healthy controls as quantified via SomaScan array (**Figure S5**). We also detected a near 4-fold increase in IGFBP2 concentrations in AVS serum relative to healthy serum samples as quantified via enzyme-linked immunosorbent assay (ELISA) (**Figure 4A**). A strong correlation between measured IGFBP2 serum concentration and IGFBP2 serum abundance was observed (Pearson’s R = 0.921), indicating accurate serum measurements from both assays (**Figure S6**). IGFBP2 concentrations were similar between male and female AVS serum samples (**Figure S7**).

**Figure 4.**
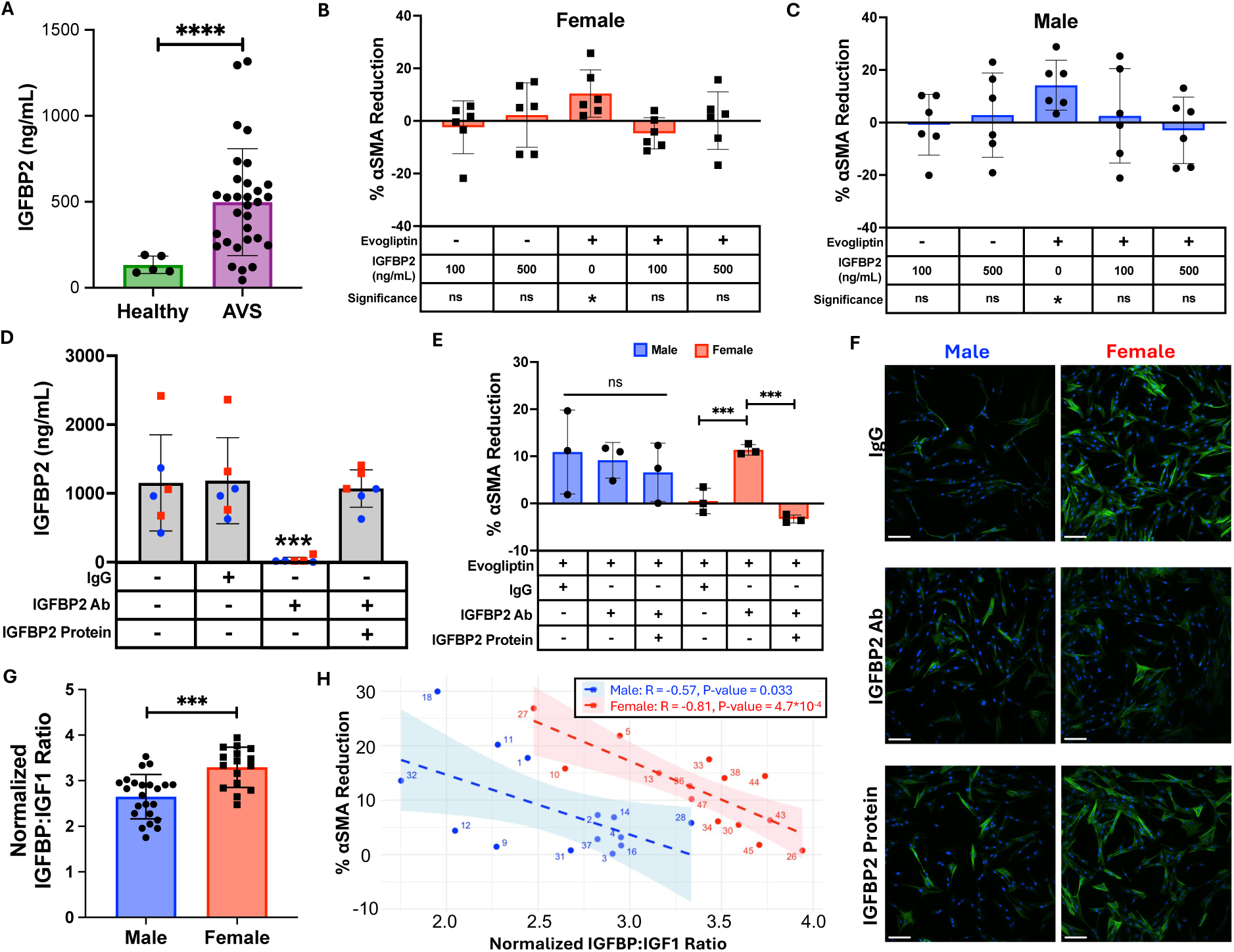
**IGFBP2 mediates Evogliptin resistance in female VICs cultured with female AVS serum**. **A,** Serum IGFBP2 concentrations quantified using an ELISA kit (N=31 AVS, N=5 healthy). Statistical significance is indicated as **** = P < 0.0001 (unpaired two-tailed t-test with Welch’s correction). **B** through **C,** Percent αSMA reduction in **(B)** female and **(C)** male VICs cultured on hydrogels with FBS and treated with Evogliptin and/or recombinant human IGFBP2 protein (N=6 gels). Statistical significance is indicated as * = P < 0.05 (one sample t and Wilcoxon test against a theoretical mean of zero). **D,** Serum IGFBP2 levels quantified using an ELISA kit with and without IGFBP2 neutralizing antibody (IGFBP2 Ab), IgG control antibody, and recombinant human IGFBP2 protein (N=3 male, N=3 female). Statistical significance is indicated as *** = P < 0.001 (one-way ANOVA with Tukey posttests). **E,** Percent αSMA reduction in male and female VICs cultured on hydrogels with sex-matched AVS patient serum samples, Evogliptin, IgG, IGFBP2 Ab, or recombinant human IGFBP2 protein (N=3 male, N=3 female). Statistical significance is indicated as *** = P < 0.001 (one-way ANOVA with Tukey posttests). **F,** Representative immunofluorescent images of male and female VICs cultured on hydrogels (green: αSMA, blue: nuclei). Scale bar = 100 μm. **G,** Normalized IGFBP to IGF1 ratio for male and female AVS serum samples (N=23 male, N=16 female). Statistical significance is indicated as *** = P < 0.001 (unpaired two-tailed t-test with Welch’s correction). **H,** Correlation between normalized IGFBP to IGF1 ratios and percent αSMA reduction in male and female VICs cultured on hydrogels with sex-matched serum and Evogliptin.

Using IGFBP2 doses corresponding to measured healthy and AVS serum levels, we next assessed the effects of IGFBP2 on Evogliptin efficacy. Female VICs cultured on hydrogels with FBS showed no baseline response to 100 or 500 ng/mL IGFBP2. However, the ability of Evogliptin to reduce myofibroblast activation was abolished in the presence of 100 or 500 ng/mL IGFBP2, confirming that physiologically relevant concentrations of IGFBP2 can regulate Evogliptin efficacy in female VICs (**Figure 4B**). Repeating this experiment with male VICs yielded similar results, suggesting that the female-specific correlation between IGFBP2 serum levels and Evogliptin efficacy is dependent on differences in serum composition rather than intrinsic male vs. female VIC drug sensitivity (**Figure 4C**).

To test this hypothesis, we next aimed to neutralize IGFBP2 in serum using an antibody and rescue IGFBP2 effects by adding excess recombinant human IGFBP2 protein. We selected the three male and female AVS serum samples with the highest IGFBP2 levels and confirmed that the neutralizing antibody effectively blocked IGFBP2 (**Figure 4D**). IGFBP2 stability in serum over the two-day culture period was also verified to ensure minimal degradation (**Figure S8**). Male or female VICs were then seeded on hydrogels and cultured with Evogliptin under three conditions: (1) sex-matched serum with a control antibody, (2) sex-matched serum with IGFBP2 neutralized, or (3) sex-matched serum with excess IGFBP2 added after neutralization. Male VICs responded to Evogliptin at baseline and showed no changes in Evogliptin response with IGFBP2 neutralization or rescue (**Figure 4E-F**). In contrast, female VICs were initially unresponsive to Evogliptin but became responsive when IGFBP2 was neutralized. Restoring IGFBP2 protein levels again negated the effects of Evogliptin, demonstrating that IGFBP2 mediates Evogliptin resistance specifically in female VICs cultured with female AVS serum.

Given the unique female response to IGFBP2 neutralization, we next investigated whether overall insulin-like growth factor 1 (IGF1) signaling is altered in female AVS serum. We calculated the normalized IGFBP:IGF1 ratio in male and female AVS serum and found that female serum exhibited a 1.25-fold higher ratio, suggesting a relative reduction in factors promoting IGF1 signaling (**Figure 4G**). Correlating this normalized insulin-like growth factor binding protein (IGFBP) to IGF1 ratio with Evogliptin efficacy in VICs cultured with sex-matched AVS serum revealed a significant negative correlation in both male and female VICs (**Figure 4H**). Together, these data indicate that an elevated IGFBP to IGF1 ratio contributes to reduced Evogliptin efficacy and that female serum is particularly susceptible to IGFBP2 mediated Evogliptin resistance due to this elevated ratio.

### RNA sequencing shows that IGFBP2 acts through mechanosensing pathways to promote Evogliptin resistance in female VICs

Next, we performed RNA sequencing to investigate the pathways by which IGFBP2 promotes resistance to Evogliptin in female VICs cultured with female AVS serum. We profiled the transcriptomes of female VICs cultured under three conditions: (1) female AVS serum alone (vehicle), (2) female AVS serum with Evogliptin (Evo), and (3) female AVS serum with Evogliptin plus an IGFBP2-neutralizing antibody (EvoAb). As expected, IGFBP2 neutralization enhanced Evogliptin efficacy, as indicated by reduced gene expression of the myofibroblast marker, *ACTA2* (**Figure S9**). Principal component analysis demonstrated that the vehicle and Evo groups clustered closely together, whereas the EvoAb group was distinctly separated (**Figure 5A**). Differential gene expression analysis further supported this observation with only 35 genes found to be significantly altered between the vehicle and Evo groups (**Figure 5B**). In contrast, 469 differentially expressed genes were detected between the vehicle and EvoAb groups (**Figure 5C**). These results indicate that IGFBP2 neutralization induces a robust transcriptomic shift during Evogliptin treatment. A comparison of differentially expressed genes relative to the vehicle group revealed 12 genes commonly enriched in both the Evo and EvoAb conditions (**Figure 5D**). Directly comparing the Evo and EvoAb groups resulted in 386 genes differentially expressed, with 182 genes upregulated in the Evo group (**Figure 5E**). Analyzing the 20 genes most upregulated or downregulated genes in the EvoAb group relative to vehicle showed that the Evo group aligned closely with the vehicle group, further confirming the inefficacy of Evogliptin treatment in female VICs cultured with female AVS serum containing high levels of IGFBP2 (**Figure 5F**).

**Figure 5.**
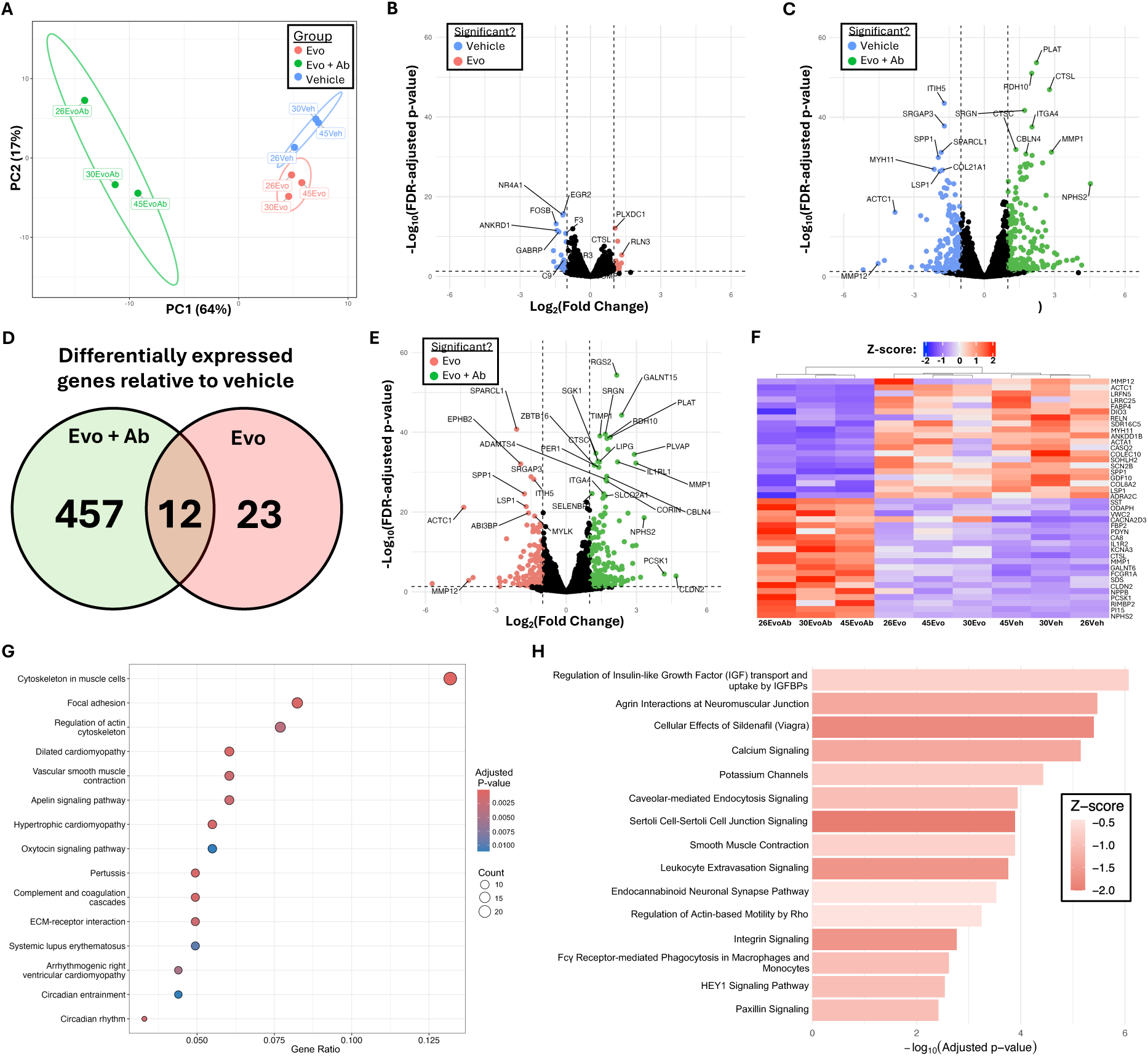
RNA sequencing identifies target pathways for IGFBP2-mediated Evogliptin resistance in female VICs cultured with female AVS serum. A, Principal component analysis clustering samples based on RNA sequencing results (N=3). **B** through **C,** Volcano plot of differential gene expression from RNA sequencing between **(B)** Vehicle and Evogliptin groups or **(C)** Vehicle and Evogliptin plus IGFBP2 neutralizing antibody groups (N=3). Statistical significance was defined as FDR-adjusted P-value < 0.05 (unpaired two-tailed t-test with Welch’s correction) and fold change greater than 2. **D,** Venn diagram comparing differentially expressed genes in the Evogliptin and Evogliptin plus IGFBP2 neutralizing antibody groups relative to vehicle control (N=3). **E,** Volcano plot of differential gene expression from RNA sequencing between Evogliptin and Evogliptin plus IGFBP2 neutralizing antibody groups (N=3). Statistical significance was defined as FDR-adjusted P-value < 0.05 (unpaired t-test with Welch’s correction) and fold change greater than 2. **F,** Heatmap of the top 20 up and down regulated genes in the Evogliptin plus IGFBP2 neutralizing antibody group relative to vehicle control (N=3). **G** through **H,** Top 15 upregulated pathways in Evogliptin versus Evogliptin plus IGFBP2 neutralizing antibody group identified by **(G)** KEGG and **(H)** Ingenuity Pathway Analysis. Abbreviations: Evo: Evogliptin, Evo + Ab: Evogliptin with IGFBP2 neutralizing antibody, Veh: Vehicle control. Number indicates patient serum sample ID (26, 30, and 45).

We next performed a pathway enrichment analysis to identify candidate signaling networks through which IGFBP2 promotes resistance to Evogliptin. Our KEGG pathway analysis on genes upregulated in the Evo group relative to the EvoAb group revealed enrichment of pathways related to mechanosensing, focal adhesion formation, and cytoskeletal regulation (**Figure 5G**). We also performed IPA and identified “Regulation of Insulin-like Growth Factor (IGF) transport and uptake by IGFBPs” as the top pathway, confirming IGFBP2 neutralization in the EvoAb condition. Additional pathways enriched in the Evo group included Rho/ROCK signaling and integrin signaling (**Figure 5H**). In contrast, the top pathways enriched in the EvoAb group relative to the Evo group were primarily associated with biochemical signaling processes (**Figure S10**). Collectively, these analyses suggest that IGFBP2 may promote Evogliptin resistance by enhancing mechanosensing and downstream integrin-mediated signaling in female VICs.

### IGFBP2 drives female-specific Evogliptin resistance through Rho/ROCK and focal adhesion kinase signaling

As a strategy to validate pathways identified through RNA sequencing, we next hypothesized that inhibiting mechanosensing pathways would sensitize female myofibroblasts to Evogliptin. Female VICs were cultured on hydrogels with female AVS patient serum and treated with inhibitors targeting Rho/ROCK or focal adhesion kinase (FAK) signaling in the presence and absence of Evogliptin (**Figure 6A**). Three doses of the Rho/ROCK inhibitor (H-1152) and the FAK inhibitor (PF-573228) were tested to identify an optimized dose that has a minimal effect on baseline myofibroblast activation but acts effectively in combination with Evogliptin (**Figure 6B-C**). We next sought to mimic the effects of IGFBP2 neutralization and assess if IGFBP2 promotes Evogliptin resistance through Rho/ROCK or FAK signaling. Using optimized doses, we demonstrated that inhibition of Rho/ROCK or FAK signaling alone has a minimal impact on female VIC myofibroblast activation. However, combining these inhibitors of mechanosensing pathways with Evogliptin restores Evogliptin efficacy in female VICs (**Figure 6D**), mirroring the effects of IGFBP2 neutralization (**Figure 4E**). Furthermore, male VICs cultured with male AVS serum were similarly responsive across all conditions tested, confirming that this mechanism is specific to female VICs cultured with female AVS serum (**Figure 6E-F**).

**Figure 6.**
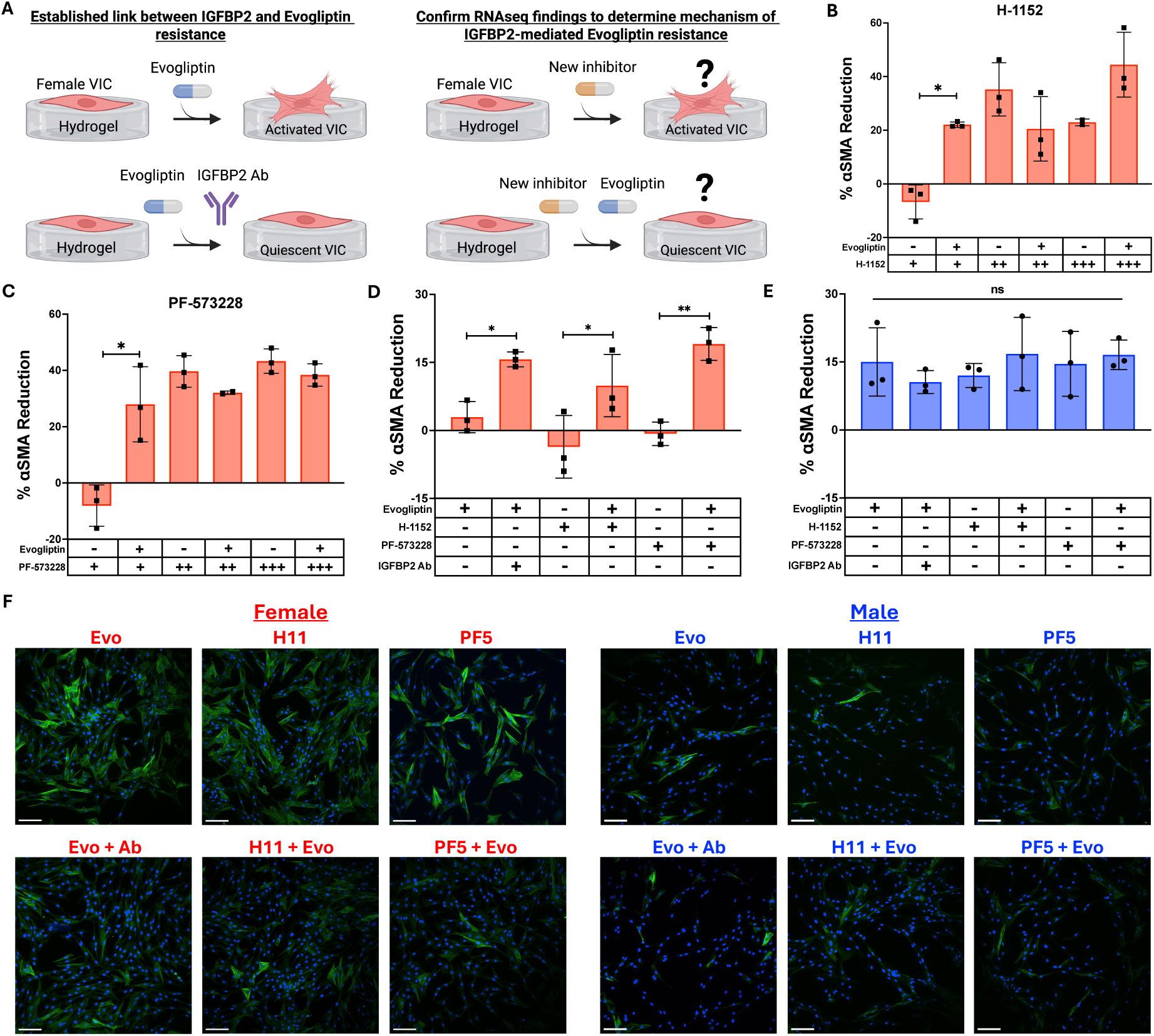
**IGFBP2 promotes Evogliptin resistance in female VICs cultured with female AVS serum through Rho/ROCK and focal adhesion kinase signaling.** A, Schematic overview of the approach to validate the mechanism of IGFBP2-mediated Evogliptin resistance. **B,** Percent αSMA reduction in female VICs cultured on hydrogels with female AVS serum and treated with various doses of H-1152 with and without Evogliptin (N=3 gels). For H-1152: + = 0.1 μM; ++ = 0.5 μM; +++ = 5 μM. Statistical significance is indicated as * = P < 0.05 (one-way ANOVA with Tukey posttests). **C,** Percent αSMA reduction in female VICs cultured on hydrogels with AVS serum and treated with various doses of PF-573228 with and without Evogliptin (N=3 gels). For PF-573228: + = 0.1 μM; ++ = 1 μM; +++ = 10 μM. Statistical significance is indicated as * = P < 0.05 (one-way ANOVA with Tukey posttests). **D** through **E,** Percent αSMA reduction in **(D)** female or **(E)** male VICs cultured on hydrogels with sex-matched AVS serum and treated with various combinations of Evogliptin, IGFBP2 neutralizing antibody, H-1152, and PF-573228 (N=3 serum samples per sex). Statistical significance is indicated as * = P < 0.05, ** = P < 0.01 (one-way ANOVA with Tukey posttests). **F,** Representative immunofluorescent images of male and female VICs cultured on hydrogels with sex-matched serum and various drug combinations (green: αSMA, blue: nuclei). Scale bar = 100 μm. Abbreviations: Evo = Evogliptin, H11 = H-1152, PF5 = PF-573228, Ab = IGFBP2 neutralizing antibody.

## Discussion

Collectively, our work revealed sex-specific associations between individual circulating serum factors and *in vitro* drug responses in male and female VICs. Although sex differences in individual serum factor abundances were minimal, the differences in correlations between serum factors and drug responses in male and female VICs were profound (**Figure 3**). This observation is consistent with prior work that identified sex-specific biomarkers that correlate with therapeutic outcomes in a multitude of other cardiovascular disease contexts.^57–59^ In our study, 19 robustly validated AVS biomarkers showed predominantly positive correlations with Ataciguat efficacy and predominantly negative correlations with Colchicine and Evogliptin efficacy in female VICs cultured with female AVS serum (**Figure S4**). In contrast, the opposite trends were observed in male VICs cultured with male AVS serum, highlighting fundamentally different biomarker and drug response relationships between sexes.

Notably, recent preliminary data from the Ataciguat clinical trial suggests that Ataciguat may be more effective at slowing calcification progression in male patients relative to female patients.^19^ In contrast, our *in vitro* data demonstrates greater efficacy of Ataciguat in reducing myofibroblast activation in female VICs cultured with female AVS serum (**Figure 3**). These differences underscore the need to better define the sex-specific mechanisms governing drug responses across experimental scales, which may arise from sex differences in differential protein binding, interactions with hormones, drug absorption, circulating metabolites, enzymatic activity, or drug pharmacokinetics.^60–62^ In contrast to Ataciguat, Colchicine and Evogliptin exhibited comparable efficacy in suppressing myofibroblast activation in male and female VICs. However, minimal data is available regarding sex-specific responses to Colchicine or Evogliptin, and no robust sex-stratified human clinical trial data is available for either of these drugs. As sex is increasingly recognized as a crucial regulator of drug efficacy, elucidating the mechanisms underlying sex-specific drug responses may provide new opportunities for optimizing precision therapies for AVS.

Of our sex-specific serum factor-drug correlations, the female-specific relationship between IGFBP2 abundance and decreased Evogliptin efficacy was our most striking finding (**Figure 3**). Evogliptin works mechanistically by inhibiting dipeptidyl peptidase 4, which increases free IGF1 and IGF1 signaling to promote fibroblast quiescence.^63,64^ IGFBP2 and other IGFBPs counteract inhibition of dipeptidyl peptidase 4 by binding to bioavailable IGF1 and inhibiting downstream IGF1 signaling.^65^ In female AVS serum, we observed an overall increased ratio of IGFBPs to IGF1 relative to male serum, suggesting lower baseline IGF1 bioavailability (**Figure 4**). Similar findings have been observed in healthy adults, where higher baseline ratios of IGFBPs to IGF1 were measured in female serum.^66^ This difference is particularly pronounced in adults over the age of 50 as menopause has been shown to further increase the ratio of IGFBPs to IGF1.^67^ We suggest that this altered balance of IGFBPs to IGF1 in female serum contributes to the observed female-specific IGFBP2-mediated resistance to Evogliptin.

Using our hydrogel cell culture platform^46^, we further demonstrate that IGFBP2 acts through Rho/ROCK and FAK signaling to desensitize female VICs to Evogliptin treatment (**Figure 6**). Aside from directly binding IGF1, IGFBP2 can also alter cellular signaling through non-canonical pathways.^68^ Notably, IGFBP2 contains an RGD motif that allows it to directly bind αvβ1 integrins, which can help stabilize focal adhesion formation and promote downstream Rho/ROCK and FAK signaling.^68–70^ Aligning with these findings, our RNA sequencing analysis identified mechanosensing pathways such as integrin signaling and focal adhesion formation among the top pathways upregulated with IGFBP2 present in female AVS serum (**Figure 5**). Moreover, mechanosensing-linked pathways such as FAK signaling have been shown to directly modulate IGF1 signaling, highlighting potential reciprocal crosstalk between these pathways.^68,71^ In our data, several of the top differentially expressed genes (*ACTA1, ACTC1, MYH11,* and *MMP12*) are well-established downstream effectors of cytoskeletal tension and extracellular matrix remodeling that are functionally connected to IGF1 activity, supporting the existence of a feedback loop between mechanosensing and IGF1 signaling. Previous work has also shown that genes that escape X chromosome inactivation intersect with the Rho/ROCK signaling pathway, revealing a potential sex chromosome contribution to female-specific mechanosensing and subsequent Evogliptin resistance.^38^ Taken together, we propose that female serum has decreased bioavailable IGF1 and excess bioavailable IGFBP2, which enhances Rho/ROCK and FAK signaling to promote resistance to Evogliptin (**Figure 7**).

**Figure 7.**
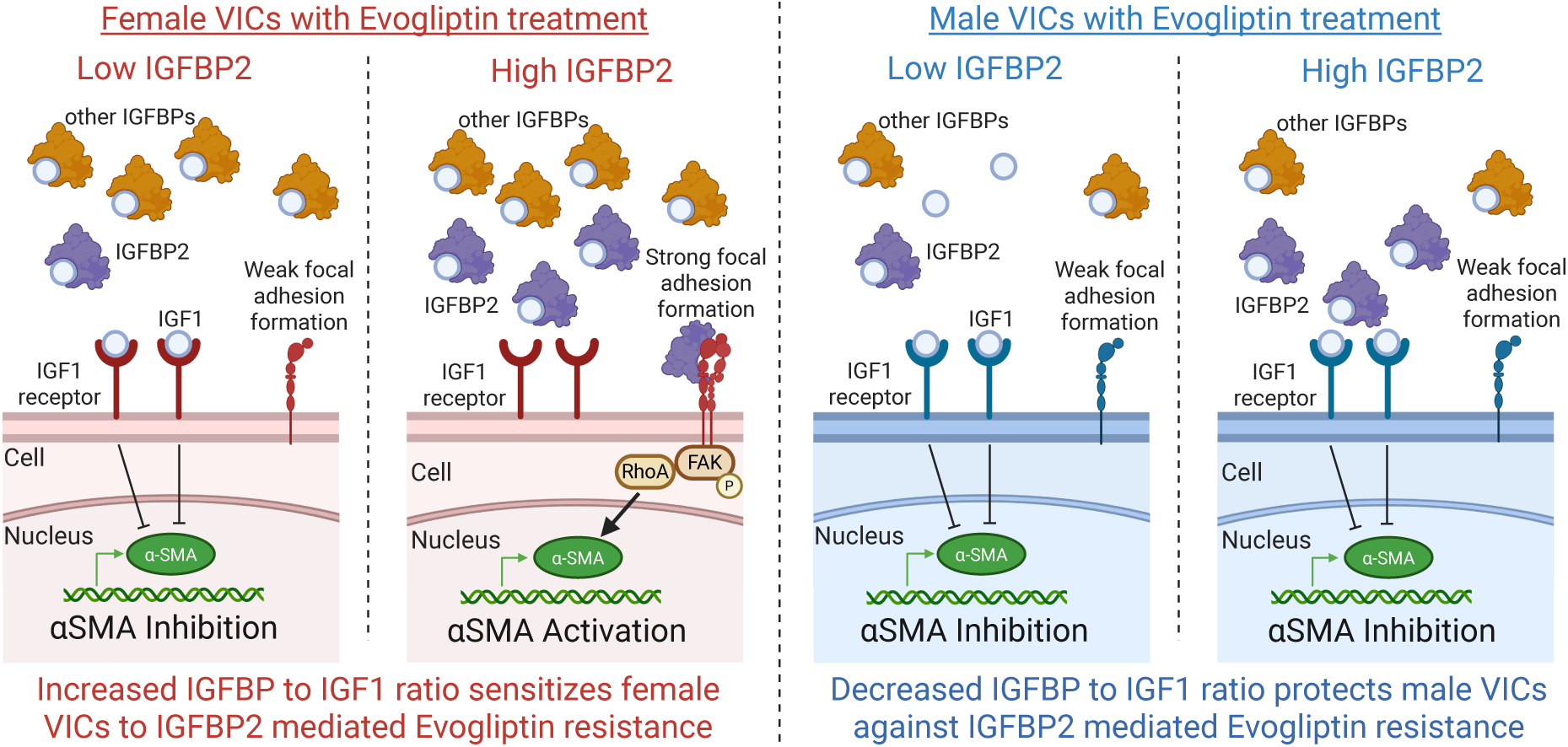
Proposed mechanism for IGFBP2 driving sex-specific Evogliptin resistance. Summary mechanistic figure illustrating how elevated IGFBP2 in female serum modulates Evogliptin efficacy in female VICs.

We also suggest that IGFBP2 may serve both as a biomarker of AVS progression and as a predictive indicator of drug responsiveness in female AVS patients. IGFBP2 has emerged as a compelling AVS biomarker due to prior clinical studies that have demonstrated its ability to predict both AVS progression and patient outcomes following transcatheter aortic valve replacement.^55,72^ Specifically, patients with circulating IGFBP2 levels above 275 ng/mL were found to be at an elevated risk of mortality following aortic valve replacement. Although no sex stratification analysis was performed, this study highlights the clinical utility of IGFBP2 as a prognostic indicator for AVS risk stratification. Our work also builds upon reports implicating IGFBP2 in altered drug sensitivity and treatment outcomes in multiple disease contexts by extending the relevance of IGFBP2 to predicting antifibrotic drug outcomes in AVS.^73–75^ A recent large-scale proteomic analysis revealed a sex-specific association between IGFBP2 abundance and AVS progression, with elevated IGFBP2 correlating with disease severity only in female patients.^76^ These findings suggest a female-specific role for IGFBP2 in predicting treatment response in AVS and underscore the importance of considering biological sex when interpreting circulating biomarkers in AVS and other cardiovascular diseases.^77^

Our study relied on the implementation of hydrogel biomaterials as VIC culture platforms to identify a female VIC-specific association between serum IGFBP2 levels and Evogliptin efficacy. Indeed, prior work has demonstrated that culturing porcine VICs with human AVS serum on hydrogels may serve as a bridge between *in vitro* disease modeling and *in vivo* patient outcomes.^38^ Future studies incorporating human-derived primary or induced pluripotent stem cell-derived VICs will be important to continue enhancing the clinical translatability of our findings.^78^ Beyond cell-specific mechanisms, the effects of organism-wide biological factors, including drug metabolism, hormone cycling, and complex physiological responses are also likely to contribute to sex-specific biomarker drug predictability.^79,80^ In addition, predicting drug responsiveness in AVS may be improved by integrating panels of multiple serum factors, such as an expanded profiling of global IGF1 signaling.^81,82^ Moreover, investigating the effects of therapeutics on VIC calcification using three-dimensional hydrogel culture systems can provide mechanistic insights into how inhibitors modulate osteoblast-like activation in male and female VICs.^44,83^ In sum, refinement of experimental models and biomarker frameworks may build upon our findings and provide additional context for sex-specific AVS patient treatment recommendations.

## Methods

### Human serum collection and processing

Patient serum samples were obtained under the approval of the University of California San Diego Institutional Review Board (IRB #804209). AVS patients were seen in a multidisciplinary valve clinic and underwent evaluation by an interventional cardiologist using echocardiography. We collected 42 AVS patient serum samples (N=25 male, N=17 female) and 5 age-matched healthy serum samples (N=3 male, N=2 female). All patients signed an IRB-approved informed consent form, and all samples were de-identified.

Collected serum samples were allowed to sit at room temperature for 15-30 minutes to allow sufficient blood clotting. Serum samples were then centrifuged at 4°C and 2,000g for 10 minutes, placed on ice, and aliquoted for storage at-80°C for up to 3 years. Serum samples were discarded after two thaw cycles.

### SomaScan proteomic analysis

All 47 serum samples were submitted for proteomic analysis using the SomaScan DNA aptamer array following protocols outlined previously.^53,54^ An additional 38 age-matched healthy serum samples (N=16 male, N=22 female) were used from the public SomaScan repository to supplement our analyses. A serum comparison analysis pipeline was developed in R to quantify patient differences in serum factor abundance. Briefly, the raw SomaScan data was trimmed and filtered to remove strong outliers (defined as quartile 1 – 3 * interquartile range and quartile 3 + 3 * interquartile range). A principal component analysis and unsupervised hierarchical clustering analysis were then performed to assess global differences in patient serum proteomes. Serum samples were then grouped by variables of interest (i.e. healthy vs diseased or male vs female) and compared via unpaired t-test to generate false discovery rate (FDR) adjusted p-values and fold changes for every serum factor (N=1,512). Significant proteins were defined as FDR-adjusted p-value less than 0.05 and a fold change greater than 1.2.

### Hydrogel formation

Hydrogels were generated in high throughput as described previously.^46^ Briefly, 96-well glass bottom well plates (Cellvis, Cat. No. P96-1.5H-N) were placed in a lightly sealed autoclave jar with an open scintillation vial containing 100 µL of mercaptopropyltrimethoxysilane (Sigma-Aldrich, Cat. No. 175617) and heated in an oven at 60°C for a minimum of 6 hours. Next, hydrogel precursor solution was prepared using phosphate buffered saline (PBS, Invitrogen, Cat. No. 14190-250), 7% wt/vol 8-arm 20 kDa polyethylene glycol-norbornene (PEG-Nb)^56^, 5 kDa PEG-dithiol (Jenkem, Cat. No. A4075-5), CRGDS cell binding motif (Bachem), and lithium phenyl-2,4,6-trimethyl-benzoylphosphinate (Fisher Scientific, Cat. No. 900889). 7 µL of hydrogel precursor solution was added directly into the center of each well in the 96-well glass bottom well plate using an EPMotion 5073l automated liquid handling system (Eppendorf). Using a custom 3D-printed stamp with glass rods, the hydrogel precursor solution was stamped flat at a height of 250 µm, inverted, and cured under 10 mW/cm^2^ of ultraviolet (UV) light for 3 minutes. The gel stamp was then removed, and newly formed hydrogels were washed with 5 % isopropyl alcohol in PBS for 20 minutes. Next, hydrogels were washed twice with PBS then incubated in VIC culture media (Media199 (Life Technologies, Cat. No. 11-043-023), 1% fetal bovine serum (FBS; Life Technologies, Cat. No. 16000069) or 1% AVS patient serum, 1 μg mL amphotericin B (Thermo Fisher, Cat. No. 15290026), 50 U/mL penicillin and 50 μg/mL streptomycin (Sigma, Cat. No. P4458)) overnight at 37°C for experimental use the following day.

For RNA sequencing and RT-qPCR experiments, 65 µL of gel precursor solution was pipetted onto a Sigmacote (Sigma) treated glass slide, then covered with a thiolated 25-mm circular glass coverslip and cured under 10 mW/cm^2^ of ultraviolet (UV) light for 3 minutes. Gels were then removed from the glass slide, placed in an untreated 6-well plate (Fisher, Cat. No. 08-772-49) and sterilized following the same wash protocol outlined above.

### VIC isolation

Biological replicates are defined as a pooled sample of VICs isolated from a minimum of three sex-matched porcine hearts. All VICs were isolated from hearts sourced from 6 to 8-month-old male and female adult pigs (Midwest Research Swine) as described previously.^84^ To summarize, aortic valve leaflets were isolated, pooled by sex, and submerged into Earle’s Balanced Salt Solution (EBSS, Sigma-Aldrich) supplemented with 1 μg/mL amphotericin B (Thermo Fisher), 50 U/mL penicillin and 50 μg/mL streptomycin (Sigma, Cat. No. P4458). After all leaflets were isolated, fresh EBSS supplemented with 250 units/mL of type II collagenase (Worthington) was added and leaflets were shaken for 30 minutes at 37°C and 5% carbon dioxide. Next, leaflets were vortexed for 30 seconds then placed in fresh EBSS/collagenase solution. Following a 60-minute incubation at 37°C and 5% carbon dioxide while shaking, leaflets were vortexed for two minutes and pressed through a 100 μm cell strainer. VICs were pelleted through a 10-minute centrifugation at 200g, then resuspended in VIC growth media (Media 199 (Life Technologies), 15% FBS, 1 μg/mL amphotericin B, 50 U/mL penicillin, and 50 μg/mL streptomycin). Lastly, VICs were seeded on T75 Falcon tissue culture treated flasks (Fisher) at 37°C and 5% carbon dioxide for expansion. Fresh VIC culture media was added every 24 to 48 hours until VICs were∼80% confluent. For long term storage, VICs were frozen down overnight at-80°C in 1 mL aliquots of 50% FBS, 45% VIC growth media, and 5% DMSO then transferred to a cell dewar for long-term storage in liquid nitrogen.

### VIC culture

Frozen aliquots of passage 1 male or female VICs were thawed and immediately resuspended in VIC growth media. Cells were pelleted through a 5-minute centrifugation at 200g and resuspended in 4 mL of fresh VIC growth media. The cell suspension was then split and seeded into two wells in a Falcon tissue culture treated 6 well plate (Fisher) for expansion. Media changes were performed every 24-48 hours until VICs achieved 80% confluency. Cells were then collected by removing the spent media, rinsing with PBS for 1 minute, then incubating each well with 1 mL of 1x trypsin (Life Technologies) for 3 minutes at 37°C and 5% carbon dioxide. VIC growth media was added in equal volume to neutralize the trypsin in each well and the two wells were pooled together. Cells were again pelleted through a 5-minute centrifugation at 200g then resuspended in 1 mL of VIC culture media. To ensure even cell confluency for experiments, VICs were counted using an automated hemocytometer then seeded at 6400 cells/well (20,000 cells/cm^2^) for immunostaining experiments and 180,000 cells/well for RT-qPCR and RNA sequencing experiments.

### Immunostaining

First, VICs were fixed in a solution of 4% paraformaldehyde solution in PBS for 20 minutes. Next, VICs were permeabilized using a solution of 0.1% Triton-X-100 (Fisher) in PBS for 1 hour and blocked with 5% bovine serum albumin in PBS for 1 hour. Depending on experimental setup, immunostaining was performed using a mouse anti-αSMA primary antibody (Abcam, 1:300) and/or a rabbit anti-cleaved caspase 3 primary antibody (Abcam, 1:250 dilution) in blocking solution for one hour at room temperature. After a 10 minute wash in PBS with 0.05% Tween20 (Sigma), secondary staining was performed for 1 hour in the dark using a solution of PBS with goat anti-mouse Alexa Fluor 488 (Life Technologies, 1:300 dilution), goat anti-rabbit Alexa Fluor 647 (Life Technologies, 1:300 dilution), 4′-6-diamidino-2-phenylindole (Life Technologies, 1:500 dilution) and HCS Cell Mask Deep Orange (Life Technologies, 1:5000 dilution). Lastly, VICs were washed once with PBS for 10 minutes then stored at 4°C in fresh PBS in the dark until imaging. Cells were imaged within 1 week of immunostaining using either a Nikon Eclipse Ti2-E or a Leica Stellaris.

### Pathway enrichment analysis

A spreadsheet containing Entrez gene IDs, fold changes between groups, and FDR-adjusted p-values was uploaded into the Ingenuity pathway analysis (IPA) software (Qiagen). A core expression analysis was performed using differentially expressed proteins (FDR < 0.05 and fold change greater than 1.2) or genes (FDR < 0.05 and fold change greater than 2) and the resultant signaling pathways were analyzed. This analysis was repeated to characterize differences between healthy and AVS patient proteomes and female VICs treated with Evogliptin with and without an IGFBP2 neutralizing antibody. Kyoto Encyclopedia of Genes and Genomes (KEGG) pathway enrichment analysis was also performed on differentially abundant proteins and genes using the ClusterProfiler package in R.

### Protein and inhibitor preparation

Lyophilized proteins were stored at-20°C for up to 6 months and resuspended following manufacturer’s guidelines (R&D systems). Recombinant human insulin-like growth factor-binding protein 2 (IGFBP2, Cat. No. 674-B2-025) was aliquoted at 100 µg/mL in sterile PBS. IGFBP2 stocks were stored at-20°C for up to 3 months and discarded after two thaw cycles.

Lyophilized inhibitors were stored at-20°C for up to 1 month then resuspended following manufacturer’s guidelines. Evogliptin (MedChem Express, Cat. No. HY-117985B), Ataciguat (MedChem Express, Cat. No. HY-17500), and PF-573228 (MedChem Express, Cat. No. HY-10461) were resuspended in sterile dimethyl sulfoxide (DMSO) (Fisher, Cat. No. D2438). Colchicine (MedChem Express, Cat. No. HY-16569) and H-1152 (Bio-techne, Cat. No. 2414) were resuspended in sterile deionized water. All aliquots were stored at-80°C for up to 6 months and discarded after two thaw cycles. Evogliptin, Ataciguat, and PF-573228 were diluted prior to experimental use to ensure a final DMSO concentration below 0.3%.

### Calculating αSMA reduction and cytotoxicity

A custom MATLAB script was used to quantify single cell αSMA gradient mean intensity data from immunostained cells. Drug efficacy was then calculated as the percent reduction in single cell αSMA gradient mean intensity of the treated condition normalized to a vehicle control as shown in the equation below:

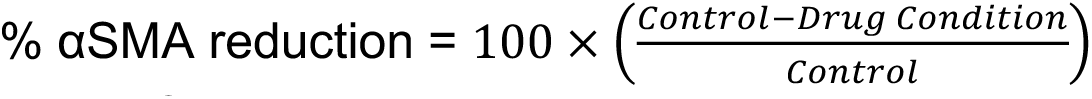

Cytotoxicity was assessed by assigning each cell in an image as alive or dead based on immunofluorescent cleaved caspase-3 expression normalized to cell mask. A positive control for cell death (10 nM Staurosporine (Cell Signaling Technologies) in VIC culture media) was used to set the cleaved caspase-3 threshold for a live/dead cell. For all αSMA and cytotoxicity data, a minimum of 100 cells were used per well with multiple wells per condition.

### Inhibitor and serum factor correlation analysis

Using R, we developed an analysis pipeline to correlate *in vitro* drug efficacy with individual serum factor abundance. Serum factor RFU values were log10 transformed and filtered to remove strong outliers as described earlier. Patient serum data was then sex-separated to perform a sex-specific correlation analysis or pooled to perform a global correlation analysis. For each analysis, the percent αSMA reduction for Ataciguat, Evogliptin, and Colchicine were aligned with individual serum factor relative fluorescent unit (RFU) values for each patient. The drug efficacy and corresponding serum factor RFU values were then plotted for each serum sample to generate a total of 31 distinct data points (16 male and 15 female). Using the corbetw2mat function in R, the correlation coefficient and corresponding p-value for each drug-serum factor combination were calculated. Significant correlations were calculated for sex-separated and combined analysis and were defined as p-value less than 0.05.

### Enzyme-linked immunosorbent assay (ELISA)

Serum concentrations of IGFBP2 were quantified using an IGFBP2 enzyme-linked immunosorbent assay (ELISA) kit (Invitrogen, Cat. No. EHIGFBP2). Following manufacturer’s guidelines, serum samples were diluted 1:500 in assay diluent A and standards were prepped using lyophilized IGFBP2 resuspended to concentrations ranging from 6000 pg/mL to 8.23 pg/mL following a three-fold dilution. 100 µL of diluted serum samples, standard curves, and controls were added to wells containing an IGFBP2 capture antibody overnight at 4°C while shaking. Wells were then washed three times with wash buffer (300 µL) then incubated with 100 µL of biotin conjugate for 1 hour while shaking. The wash step was then repeated before adding 100 µL of streptavidin-HRP solution (Invitrogen) for 45 minutes with gentle shaking. The wash step was repeated one last time, then 100 µL of TMB substrate (Invitrogen) was added for 30 minutes in the dark while shaking. Finally, 50 µL of stop solution (Invitrogen) was added and the plate was immediately imaged at 450 nm using a Perkin Elmer Envision plate reader. The data from the standards was fed into GraphPad Prism to generate a standard curve. The standard curve was then used to convert OD450 values to IGFBP2 concentrations in each serum sample, followed by multiplication with the serum dilution factor (500).

### Neutralizing antibody experiments

Human IGFBP2 neutralizing antibody (R&D systems, Cat. No AF674) and normal IgG control antibody (R&D systems, Cat. No AB-108-C) were resuspended in sterile PBS at 200 µg/mL and 1,000 µg/mL respectively. Antibodies were stored at-20°C for up to 6 months and discarded after two thaw cycles. IGFBP2 neutralizing antibody and normal IgG control antibody were added at 1 µg/mL to media containing AVS patient serum and incubated at 37°C for 1 hour to allow for neutralization prior to cell seeding.

### RT-qPCR

25 mm hydrogels seeded with VICs were inverted into cell lysis buffer (Qiagen RNEasy Micro Kit) for two minutes. The hydrogels were then flipped over and rinsed with 70% ethanol. The solution was then collected in spin columns. RNA was collected for all samples following manufacturers guidelines and tested for quality using a NanoDrop 2000 spectrophotometer (Thermo Fisher). RNA was then converted to cDNA using an iScript Synthesis kit (Bio-Rad). Relative RNA expression was quantified using IQ SYBR Green Supermix (Bio-Rad) fluorescent intensity measured on a CFX384 iCycler (Bio-Rad). Genes were normalized to the *RPL30* gene and quantified using the ΔΔCT method. Forward and reverse primer sequences can be found in **Table S6**.

### RNA sequencing

RNA was isolated for sequencing following the protocol described above and assessed for quality using an Agilent TapeStation 4200. A minimum of 100 ng of RNA per sample was submitted to The Sanford Consortium for Regenerative Medicine Genomics Core, which generated cDNA libraries and performed indexing. RNA sequences were assessed for quality using fastqc, and adapter sequences were removed using trimomatic (v0.39) if needed. Samples were then aligned to the porcine genome (SusScrofa11.1) using HISAT2 (v2.2.1) and trimmed to remove genes with counts less than 10.

Differential gene expression analysis was performed in R. Briefly, RNA samples were grouped by experimental condition and compared via unpaired t-test to generate false discovery rate (FDR) adjusted p-values and fold changes for every gene measured. Significant genes were defined as FDR-adjusted p-value less than 0.05 and a fold change greater than 2. Principal component analysis and pathway enrichment were performed as described previously.

## Statistical analysis

Unless otherwise stated, statistical analysis was performed in GraphPad Prism and significance was defined as a p-value less than 0.05. Comparisons between two groups were calculated by an unpaired t-test with Welch’s correction and comparisons between multiple groups were calculated via one-way ANOVA with Tukey posttests. All data are shown as mean ± standard deviation.

## Author contributions

B.J.V. and B.A.A. conceived and supervised the study. B.J.V., M.C., and R.M.G. performed all *in vitro* experiments. B.J.V., M.C., and R.M.G. analyzed data and performed statistical analyses. N.E.F.V., and R.R.R. collected and processed human serum samples. B.J.V., and B.A.A. wrote and edited the manuscript. All authors approved the manuscript.

## Materials and correspondence

All correspondence should be addressed to Brian Aguado.

## Acknowledgements

The authors would like to thank the patients that donated their serum to research. The authors acknowledge technical, sequencing, and research support provided by Dr. Trevor Biddle at the gCore in the Sanford Consortium for Regenerative Medicine and Dr. Kristen Jepsen at the Institute for Genomic Medicine in University of California San Diego. The authors also acknowledge image analysis assistance and code development provided by Dr. Peng Guo at the Nikon Imaging Center in University of California San Diego.

