## Supplementary Materials for "Circulating biomarkers in serum from aortic valve stenosis patients predict sex-specific drug responses in valve myofibroblasts"

This Supplementary Information includes:

- Supplementary Figures S1-10
- Supplementary Tables S1-6

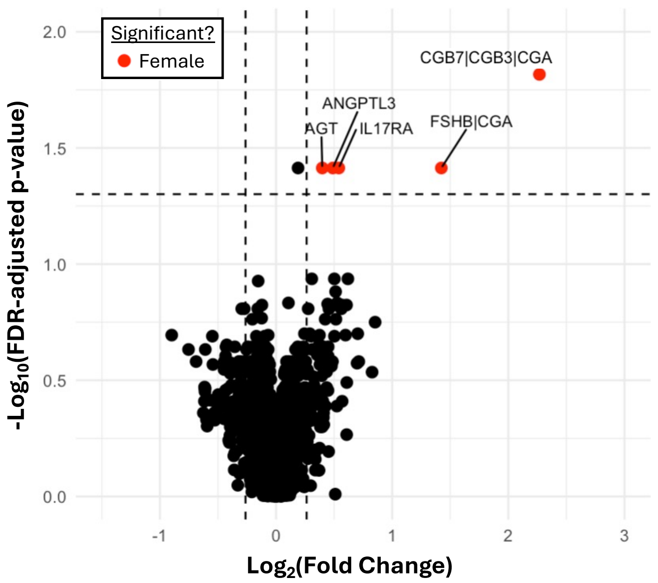


**Figure S1. Male and female AVS patient serum proteomes are similar.** Volcano plot showing proteins that are more abundant in female AVS serum samples (red) (N=1,512 proteins, N=25 male serum samples, N=17 female serum samples). Statistical significance was defined as FDR-adjusted P-value < 0.05 (unpaired two-tailed t-test with Welch's correction) and fold change greater than 1.2.

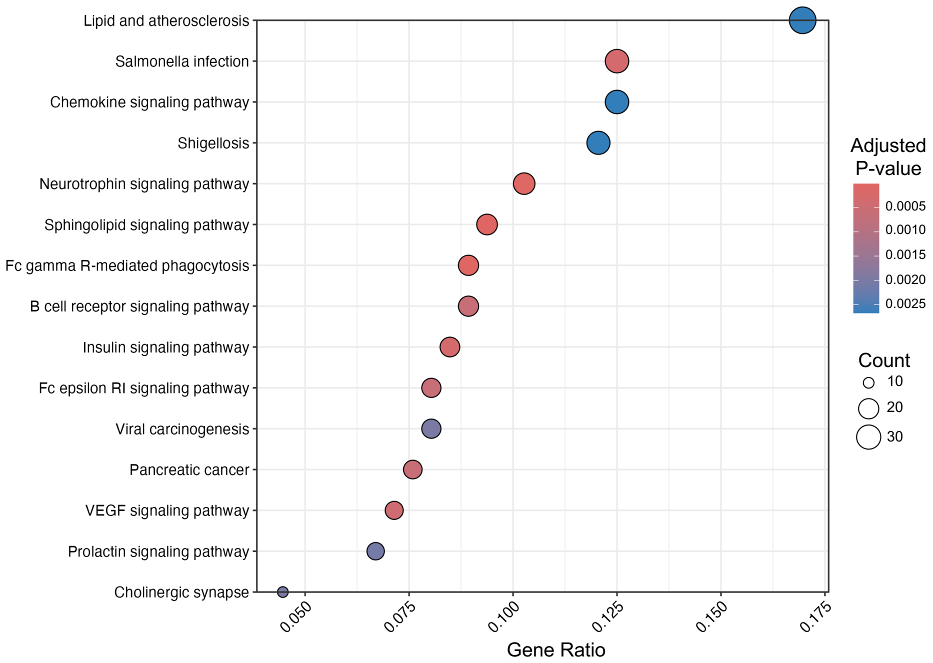


**Figure S2. Top enriched pathways identified from AVS serum samples.** Top 15 pathways identified through KEGG analysis from the proteins that are more abundant in AVS serum samples.

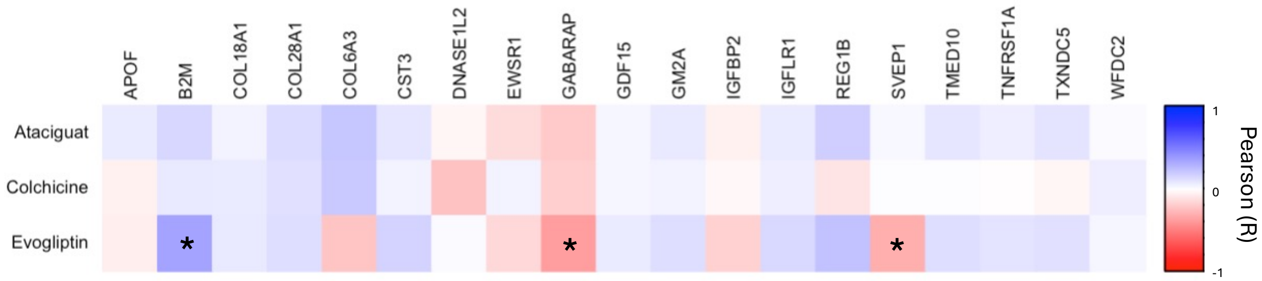


**Figure S3. Overall serum factor-drug correlations.** Heatmap showing the overall correlation between each candidate serum factor and antifibrotic drug (N=31 serum samples). Statistical significance is indicated as * = P < 0.05.

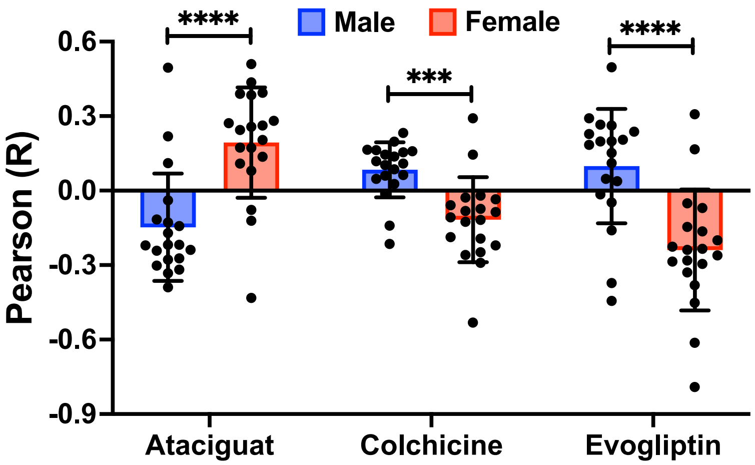


**Figure S4. Serum factor correlations with antifibrotic drugs are sex-specific**. Pearson correlation of male and female serum factors with Ataciguat, Colchicine, or Evogliptin efficacy in male and female VICs cultured on hydrogels with sex-matched AVS serum (N=19 serum factors). Statistical significance is indicated as *** = P < 0.001; **** = P < 0.0001 (unpaired two-tailed t-test with Welch's correction).

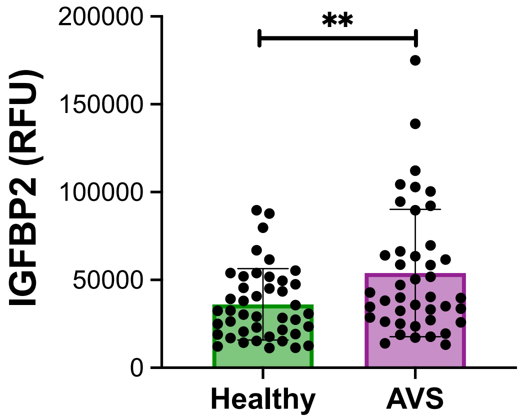


**Figure S5. IGFBP2 is more abundant in AVS patient serum samples.** Serum IGFBP2 abundance in AVS patient and healthy serum samples quantified using relative fluorescent units (RFU) (N=42 AVS, N=43 healthy). Statistical significance is defined as P-value < 0.01 (unpaired two-tailed t-test with Welch's correction).

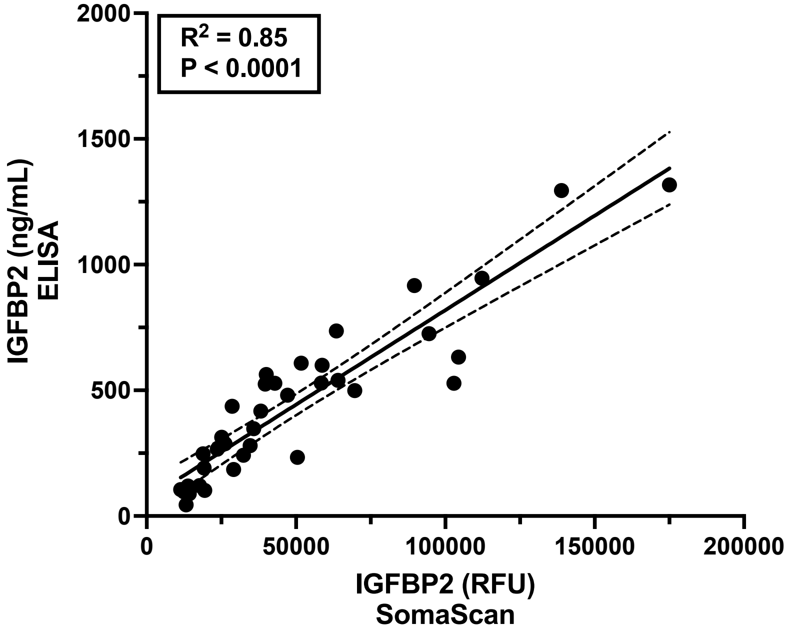


**Figure S6. IGFBP2 serum concentration is strongly correlated with IGFBP2 serum abundance.** Correlation plot comparing AVS patient serum IGFBP2 abundance quantified from the SomaScan array and serum IGFBP2 concentration quantified from an ELISA kit (N=36 serum samples).

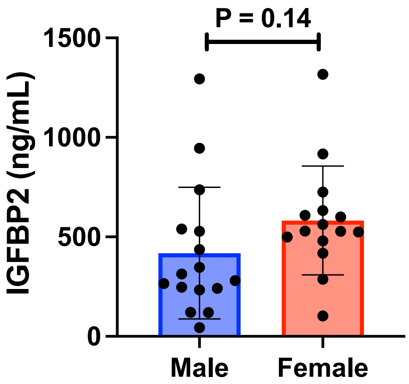


**Figure S7. Serum IGFBP2 concentration is not sex dependent.** Bar graph comparing IGFBP2 concentrations in male and female AVS serum samples (N=16 male, N=15 female). Statistical significance was determined by unpaired two-tailed t-test with Welch's correction.

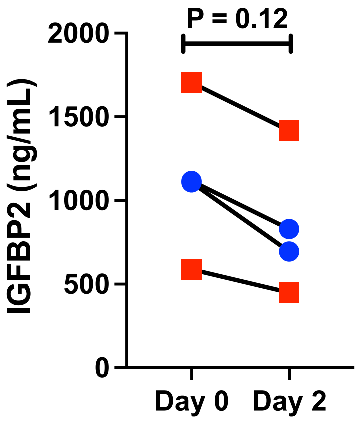


**Figure S8. IGFBP2 degradation in cell culture is minimal.** Serum IGFBP2 levels after 0 days and 2 days of culture with sex-matched male (blue) and female (red) VICs on hydrogels (N=2 serum samples per sex). Statistical significance was determined by Wilcoxon test.

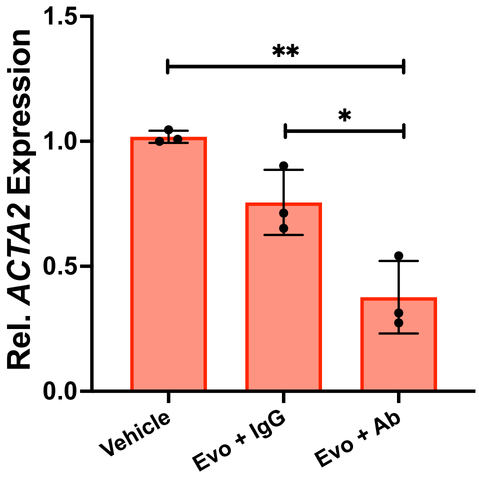


**Figure S9. ACTA2 gene expression is more robustly reduced by combining Evogliptin and IGFBP2 neutralizing antibody treatment.** RT-qPCR of *ACTA2* gene expression in vehicle, Evogliptin plus IgG antibody (Evo + IgG), and Evogliptin plus IGFBP2 neutralizing antibody (Evo + Ab) treated groups (N=3 serum samples). Statistical significance is indicated as * = P <0.05 and ** = P < 0.01 (one-way ANOVA with Tukey posttests).

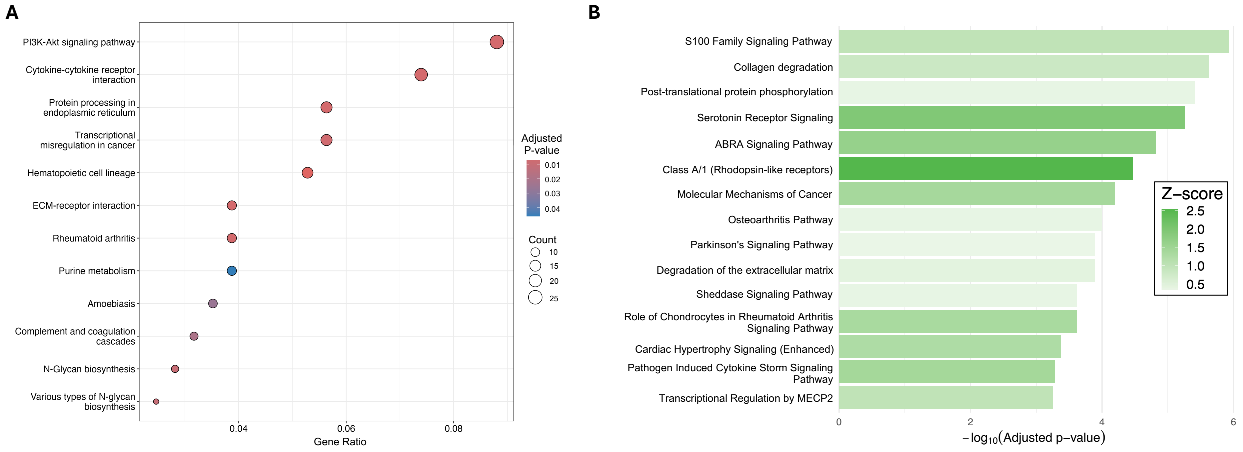


**Figure S10. Enriched pathways in female VICs treated with IGFBP2 neutralizing antibody and Evogliptin. A** through **B,** Top 15 upregulated pathways in the Evogliptin plus IGFBP2 neutralizing antibody group versus the Evogliptin group, identified by **(A)** KEGG and **(B)** Ingenuity Pathway Analysis.

**Table S1.** Male and female AVS patient clinical data comparisons. Statistical significance was determined by unpaired t-test with Welch's correction. Bolded p-values are significant (P<0.05).

| Characteristic | Male Cohort (N = 25) | Female Cohort (N = 17) | P-value |
|---|---|---|---|
| Age (years) | 78 ± 8 | 84 ± 6 | **0.016** |
| Aortic valve max velocity (m/s) | 3.91 ± 0.97 | 4.00 ± 0.52 | 0.702 |
| Aortic valve area ($m^2$) | 1.15 ± 0.85 | 0.76 ± 0.18 | **0.035** |
| Mean Pressure Gradient (mmHg) | 37.4 ± 18.2 | 38.1 ± 9.0 | 0.867 |
| SVI ($mL/m^2$) | 52.4 ± 20.9 | 46.3 ± 14.1 | 0.270 |
| LVIDD (cm) | 4.44 ± 0.77 | 3.96 ± 0.79 | 0.063 |
| LVIDS (cm) | 2.85 ± 0.77 | 2.63 ± 0.77 | 0.364 |
| Ejection fraction (%) | 62.2 ± 11.0 | 64.4 ± 11.3 | 0.540 |
| WBC ($1000/mm^2$) | 7.39 ± 2.17 | 7.32 ± 1.87 | 0.911 |
| Diastolic Volume (ml) | 96.3 ± 33.2 | 77.4 ± 39.5 | 0.134 |
| Systolic Volume (ml) | 33.2 ± 13.4 | 35.9 ± 31.6 | 0.752 |
| Aspirin (%) | 60 | 53 | 0.661 |
| Statin (%) | 17 | 25 | 0.585 |
| Losartan (%) | 64 | 65 | 0.964 |

**Table S2.** Clinically relevant dosing parameters for Ataciguat, Colchicine, and Evogliptin.

| Drug | 5% Cmax (μM) | 10% Cmax (μM) | 50% Cmax (μM) |
|---|---|---|---|
| Ataciguat | 3.47 | 6.94 | 34.7 |
| Colchicine | $7.76*10^{-4}$ | $1.55*10^{-3}$ | $7.76*10^{-3}$ |
| Evogliptin | $2.70*10^{-3}$ | $5.41*10^{-3}$ | $2.70*10^{-2}$ |

**Table S3.** Overall correlations (Pearson's R) between candidate serum biomarker abundance and percent αSMA reduction in male and female VICs cultured on hydrogels with sex-matched AVS serum and antifibrotic drugs (N=42 serum samples). Bolded correlations are significant (P<0.05).

| Protein | Overall Correlation (Pearson's R) with Ataciguat | Overall Correlation (Pearson's R) with Colchicine | Overall Correlation (Pearson's R) with Evogliptin |
|---|---|---|---|
| APOF | 0.072 | -0.055 | -0.066 |
| B2M | 0.153 | 0.085 | **0.359** |
| COL18A1 | 0.045 | 0.080 | 0.081 |
| COL28A1 | 0.132 | 0.111 | 0.125 |
| COL6A3 | 0.220 | 0.210 | -0.229 |
| CST3 | 0.095 | 0.050 | 0.164 |
| DNASE1L2 | -0.037 | -0.238 | 0.014 |
| EWSR1 | -0.137 | 0.043 | -0.146 |
| GABARAP | -0.208 | -0.188 | **-0.378** |
| GDF15 | 0.033 | 0.037 | 0.073 |
| GM2A | 0.082 | 0.040 | 0.129 |
| IGFBP2 | -0.054 | -0.020 | -0.177 |
| IGFLR1 | 0.076 | 0.066 | 0.143 |
| REG1B | 0.181 | -0.110 | 0.236 |
| SVEP1 | 0.027 | 0.004 | **-0.312** |
| TMED10 | 0.093 | 0.004 | 0.123 |
| TNFRSF1A | 0.062 | -0.009 | 0.102 |
| TXNDC5 | 0.106 | -0.033 | 0.116 |
| WFC2 | 0.019 | 0.069 | 0.032 |

**Table S4.** Correlations (Pearson's R) between male candidate serum biomarker abundance and percent αSMA reduction in male VICs cultured on hydrogels with serum and antifibrotic drugs (N=25 serum samples). Bolded correlations are significant ($P<0.05$).

| Protein | Male Correlation (Pearson's R) with Ataciguat | Male Correlation (Pearson's R) with Colchicine | Male Correlation (Pearson's R) with Evogliptin |
|---|---|---|---|
| APOF | -0.318 | 0.047 | 0.038 |
| B2M | 0.219 | 0.137 | **0.497** |
| COL18A1 | -0.273 | 0.154 | 0.205 |
| COL28A1 | -0.039 | 0.232 | 0.228 |
| COL6A3 | **0.495** | 0.165 | -0.160 |
| CST3 | -0.218 | 0.145 | 0.266 |
| DNASE1L2 | 0.111 | -0.141 | -0.048 |
| EWSR1 | -0.302 | 0.197 | 0.047 |
| GABARAP | -0.171 | -0.215 | -0.372 |
| GDF15 | -0.218 | 0.103 | 0.198 |
| GM2A | -0.242 | 0.086 | 0.199 |
| IGFBP2 | -0.333 | 0.027 | -0.015 |
| IGFLR1 | -0.239 | 0.159 | 0.262 |
| REG1B | -0.128 | -0.013 | 0.291 |
| SVEP1 | -0.221 | 0.109 | **-0.444** |
| TMED10 | -0.143 | 0.060 | 0.237 |
| TNFRSF1A | -0.278 | 0.118 | 0.184 |
| TXNDC5 | -0.116 | 0.063 | 0.152 |
| WFC2 | -0.389 | 0.163 | 0.111 |

**Table S5.** Correlations (Pearson's R) between female candidate serum biomarker abundance and percent αSMA reduction in female VICs cultured on hydrogels with serum and antifibrotic drugs (N=17 serum samples). Bolded correlations are significant ($P<0.05$).

| Protein | Female Correlation (Pearson's R) with Ataciguat | Female Correlation (Pearson's R) with Colchicine | Female Correlation (Pearson's R) with Evogliptin |
|---|---|---|---|
| APOF | 0.385 | -0.193 | -0.329 |
| B2M | -0.432 | 0.145 | 0.166 |
| COL18A1 | 0.136 | -0.020 | **-0.613** |
| COL28A1 | 0.262 | -0.126 | -0.226 |
| COL6A3 | -0.078 | 0.291 | -0.381 |
| CST3 | 0.173 | -0.086 | -0.261 |
| DNASE1L2 | 0.272 | **-0.531** | 0.308 |
| EWSR1 | 0.257 | -0.259 | -0.452 |
| GABARAP | 0.109 | -0.221 | -0.295 |
| GDF15 | 0.080 | -0.035 | -0.281 |
| GM2A | **0.510** | -0.028 | -0.146 |
| IGFBP2 | -0.121 | -0.059 | **-0.791** |
| IGFLR1 | 0.436 | -0.108 | -0.234 |
| REG1B | 0.244 | -0.291 | -0.070 |
| SVEP1 | 0.172 | -0.118 | -0.200 |
| TMED10 | 0.281 | -0.073 | -0.239 |
| TNFRSF1A | 0.394 | -0.248 | -0.164 |
| TXNDC5 | 0.204 | -0.187 | -0.051 |
| WFC2 | 0.390 | -0.082 | -0.286 |

**Table S6.** Forward and reverse primer sequences for genes assessed via RT-qPCR. All primers are shown from the 5' to 3' end.

| Gene | Forward Primer (5'-3') | Reverse Primer (5'-3') |
|---|---|---|
| *RPL30* | AGATTTCCTCAAGGCTGGGC | GCTGGGGTACAAGCAGACTC |
| *ACTA2* | GCAAACAGGAATACGATGAAGCC | AACACATAGGTAACGAGTCAGAGC |
